# Neural oscillations track natural but not artificial fast speech: Novel insights from speech-brain coupling using MEG

**DOI:** 10.1101/2020.10.20.344895

**Authors:** Ana Sofía Hincapié Casas, Tarek Lajnef, Annalisa Pascarella, Hélène Guiraud, Hannu Laaksonen, Dimitri Bayle, Karim Jerbi, Véronique Boulenger

## Abstract

Speech processing is supported by the synchronization of cortical oscillations to its rhythmic components, including syllable rate. This has been shown to be the case for normal rate speech as well as artificially accelerated speech. However, the case of natural speech rate variations, which are among the most ubiquitous sources of variability in speech, has been largely overlooked. Here, we directly compared changes in the properties of cortico-acoustic coupling when speech naturally shifts from normal to fast rate and when it is artificially accelerated. Neuromagnetic brain signals of 24 normal-hearing adults were recorded with magnetoencephalography (MEG) while they listened to natural normal (∼6 syllables/s), natural fast (∼9 syllables/s) and time-compressed (∼9 syllables/s) sentences, as well as to envelope-matched amplitude-modulated noise. We estimated coherence between the envelope of the acoustic input and MEG source time-series at frequencies corresponding to the mean syllable rates of the normal and fast speech stimuli. We found that listening to natural speech at normal and fast rates was associated with coupling between speech signal envelope and neural oscillations in right auditory and (pre)motor cortices. This oscillatory alignment occurred at ∼6.25 Hz for normal rate sentences and shifted up to ∼8.75 Hz for naturally-produced fast speech, mirroring the increase in syllable rate between the two conditions. Unexpectedly, despite being generated at the same rate as naturally-produced fast speech, the time-compressed sentences did not lead to significant cortico-acoustic coupling at ∼8.75 Hz. Interestingly, neural activity in putative right articulatory cortex exhibited stronger tuning to natural fast rather than to artificially accelerated speech, as well as stronger phase-coupling with left temporo-parietal and motor regions. This may reflect enhanced tracking of articulatory features of naturally-produced speech. Altogether, our findings provide new insights into the oscillatory brain signature underlying the perception of natural speech at different rates and highlight the importance of using naturally-produced speech when probing the dynamics of brain-to-speech coupling.

## Introduction

Understanding the neural mechanisms at play during natural spoken language comprehension remains a challenging issue, especially given the large variability of the speech acoustic signal in everyday life. For communication to be efficient, the brain has to rapidly accomplish a cascade of sophisticated operations. One of the first is to parse the continuous acoustic stream into smaller units that will then be mapped onto internal linguistic representations. In this regard, the (quasi-)rhythmicity of speech is fundamental as it allows the listener’s cognitive system to make predictions about the incoming signal, thus helping speech segmentation and comprehension (1,2). Neurocognitive models of speech perception assign a key functional role to ongoing neural oscillations in the tracking of speech rhythm (3–7). By aligning to speech at multiple timescales, brain oscillatory activity in the gamma (∼25-40 Hz), theta (∼4-7 Hz) and delta (∼1-3 Hz) frequency bands would segment the acoustic stream into phoneme-, syllable- and word-sized packets, respectively. These units may then be integrated hierarchically for higher-order linguistic processes. Convincing evidence from electro- and magnetoencephalography (EEG/MEG) revealed coupling between theta-band oscillations in the auditory cortex and the slow modulations (2-8 Hz) in speech amplitude envelope (8–13). Modulations in this range are inherently tied to syllable production (14) and are crucial for speech intelligibility as they convey, among other information, prosodic cues such as stress and tempo (15). Interestingly, auditory cortex oscillations are therefore able to track and align to the speech input in a frequency range which coincides with the average syllable rate of speakers across languages (16).

Speech rate can however substantially vary within and between speakers and contexts. As listeners, we need to rapidly adapt to the changing rates for efficient understanding (17,18). Surprisingly, only a few studies so far examined brain-to-speech coupling in the case of speech rate variations, and we know little about the spatial and frequency dynamics of cortical oscillations involved in processing naturally accelerated speech. Most of previous studies used time-compressed speech, where the duration is artificially reduced but the spectral content is kept intact (19–22). Results showed that brain-to-speech coupling occurs for moderately time-compressed, still intelligible speech but not for higher compression rates, yielding unintelligible stimuli (23). Accordingly, it has been suggested that the efficiency of speech decoding may depend on the capacity of neural oscillations (primarily in the theta range) to remain in sync with the syllable rate. Once the latter exceeds the upper limit of the neural theta band, comprehension has been found to deteriorate (1,8,24–26). This said, one EEG study (27) reported cortical coupling to the syllabic structure of time-compressed speech up to 14 Hz, even for poorly understood sentences (i.e. > 10 syllable/s). This suggests that neural oscillations are able to align to the incoming speech signal at higher frequencies to match its temporal structure, at least for artificial acceleration (see also 30) and even if speech is not fully intelligible (but see 31 for convincing evidence of a functional contribution of theta-band cortical tracking to speech intelligibility).

Taken together, previous studies of neural coupling to fast rate speech have mainly focused on artificially compressed stimuli and have led to heterogenous and partly contradictory results. In fact, although insightful, studying the perception of time-compressed speech may not provide the best model of how neural oscillations handle natural changes in speech rate. This question has been largely overlooked in the literature, yet it seems crucial given the subtle differences between naturally-produced fast speech and artificially accelerated speech. Artificial and natural acceleration of speech rate both reduce the acoustic signal in terms of length of acoustic cues, formant transitions and pauses. However, and by contrast to time-compressed speech, these changes operate non-linearly when we naturally speak at a fast rate, partly due to articulatory restrictions (30,31). In French and English for instance, vowel duration and unstressed syllables (for English) are more reduced than consonant duration and stressed syllables. Besides temporal reduction, natural fast speech also undergoes a series of spectro-temporal changes resulting in increased processing load for the listener as compared to time-compressed speech (30,32,33). Uttering speech at a fast rate decreases the spatial magnitude of articulatory movements (i.e., they are achieved more quickly and less accurately) and enhances coarticulation (i.e., increased gestural overlap) and assimilation, which can even lead to the suppression of whole segments (34). Accordingly, the listener’s auditory system faces a major challenge, namely to adjust not only to a shortened (as in time-compressed speech) but also spectro-temporally degraded signal for efficient decoding. Although adaptation to naturally accelerated speech has been reported behaviourally (17), the underlying brain oscillatory dynamics remain, to our knowledge, largely underinvestigated. In a compelling MEG study, Alexandrou and colleagues (35) recently reported alignment of auditory and parietal cortex oscillations to speech spontaneously produced at different rates (from ∼2 to 7 syllables/s). Yet their fast rate condition falls within the canonical theta range (4-7 Hz), and therefore does not address the question whether neural oscillations change their coupling frequency beyond the theta limit to track natural speech acceleration. More generally, and to the best of our knowledge, no study to date has directly compared neural entrainment to speech using either natural fast rate speech or artificially accelerated stimuli. Such a comparison may elucidate whether using artificial acceleration of speech stimuli accurately captures the brain mechanisms at play during perception of natural speech rate changes.

Here we address this question using an unprecedented MEG experiment where we compare the modulations of cortico-acoustic tracking patterns induced by normal and fast speech, generated either naturally or using time compression. Although seemingly subtle, the distinction we address is of fundamental importance to better understand how our brains track and encode the spectro-temporal changes we encounter in daily communication. Furthermore, observing differences in the neural processing of naturally versus artificially accelerated speech could be key to improving oscillatory models of speech perception. Based on previous work (27,28), one would expect that increases in speech rate would be associated with upward shifts in cortico-acoustic coupling frequency, matching the change in syllable rate, and that this would occur both for natural and artificial acceleration of the stimuli. However, given the articulatory changes elicited by natural acceleration (34) and the tight relationship between speech perception and production processes (36–38), the parsing process may particularly engage sensorimotor mechanisms during natural fast speech perception. As a result, we expect neural coupling to occur not only in auditory but also in motor regions and importantly, that this motor resonance may be stronger for naturally accelerated than for artificially manipulated speech.

To test these assumptions, we sought to unravel the oscillatory brain signature of speech naturally produced at a normal or fast rate, and of time-compressed speech. We investigated whether neural oscillations in auditory and motor cortex, recorded with MEG, align to syllable rate when it is naturally accelerated and how this compares to artificially manipulated speech. To this end, we computed cortico-acoustic coherence at the source level while participants listened to single sentences naturally produced at a normal and fast rate, or time-compressed. Most previous work assessed neural coupling to speech amplitude envelope in the canonical theta band (e.g., 9,12,34), yet speakers can naturally slow down or speed up their syllable rate outside the limits of this range. Examining cortical tracking of speech rate variations in frequency ranges that specifically match the temporal structure of heard speech therefore appears as a more straightforward approach (39). Accordingly, we assessed brain synchronization to the envelope of normal and fast rate speech at two frequencies of interest, identified as peaks in the respective power spectra of the speech signals. By including amplitude-modulated noise control stimuli (based on the temporal envelopes of normal and fast rate sentences), we additionally investigated whether cortico-acoustic coupling depends on the presence of linguistic content in the stimuli or simply reflects brain responses to low-level acoustic cues. Finally, the results of the speech-brain coupling were used to further examine the oscillatory network dynamics using seed-based phase-coupling analyses.

## Results

We recorded MEG brain activity of 24 healthy French adult participants during perception of a set of natural normal rate (mean = 6.76 ± 0.57 syllables/s), natural fast rate (9.15 ± 0.60 syllables/s) and time-compressed sentences (to the same syllable rate as natural fast rate speech). Amplitude-modulated noise stimuli (modulated with normal rate and fast rate sentence envelopes) served as non-speech control conditions. Participants were instructed to attentively listen to the different stimuli and detect beep-sounds embedded in filler sentences (not analyzed) by pressing a button as quickly as possible. To assess neural tracking of speech rate variations, we computed cortico-acoustic coherence between signal’s amplitude envelope and source-localized MEG time-series (see Fig. 1 for an overview of the method). We defined two frequencies of interest for analysis (∼6.25Hz and ∼8.75 Hz), based on the frequency peaks identified in the power spectra of normal and fast rate sentences respectively (Fig. 2, see also supplementary Fig. S1 and Table S1). These peaks closely matched the mean syllable rate of the speech stimuli, derived by assessing the number of syllables over time with Praat (40). For statistical analysis, we contrasted coherence measures obtained for actual stimulus encoding with coherence values obtained using surrogate data, as well as with during a pre-stimulus baseline. We generated the surrogate data by randomly shuffling the speech trial order so that they no longer matched the associated MEG signals (see Material and Methods).

**Fig 1.**
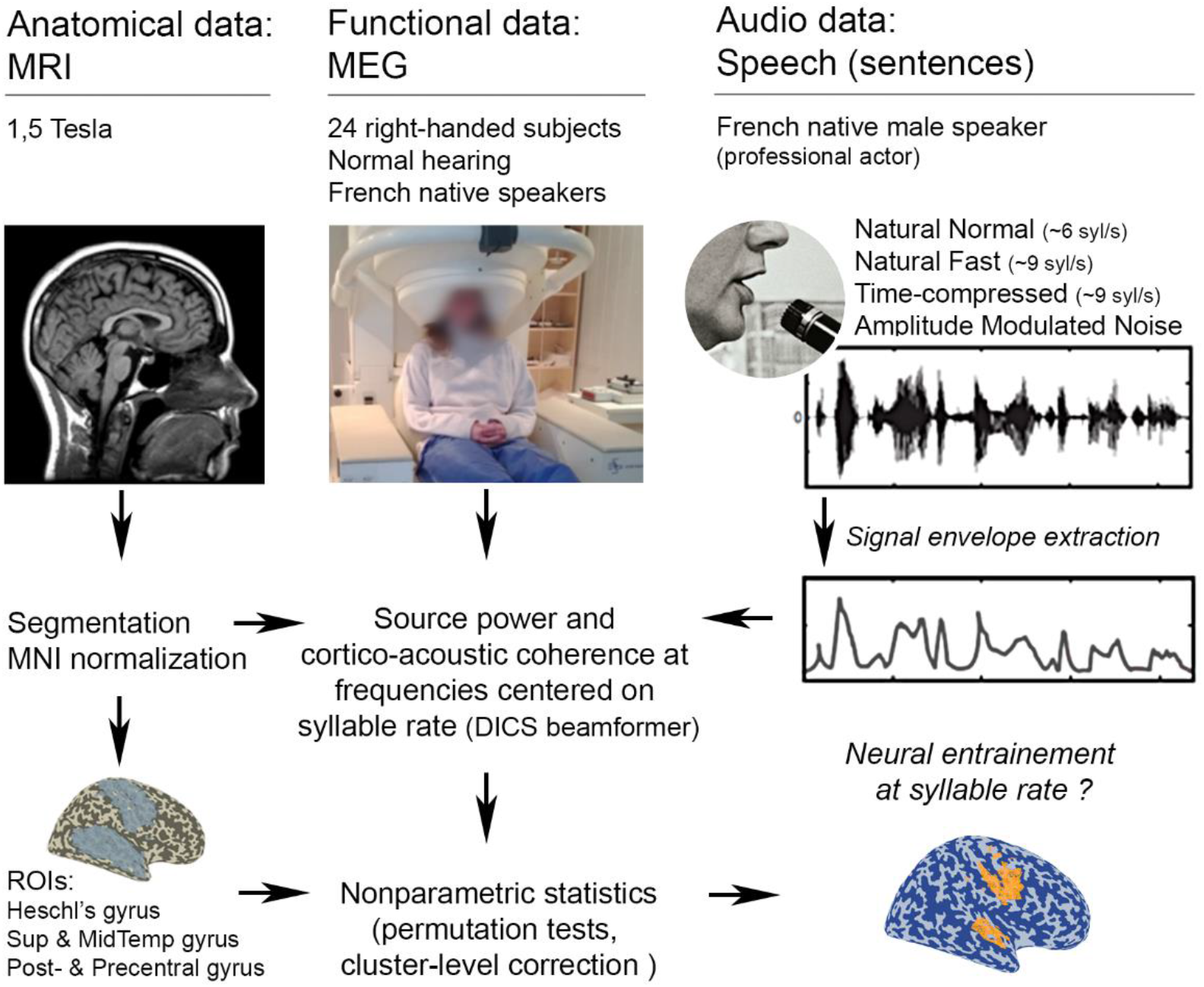
Overview of the cortico-acoustic coupling analysis. Computation of cortico-acoustic coherence required three inputs: the reference signal, namely the audio recordings; the anatomical data, i.e. the participants’ MRI; and the functional data, i.e. MEG recordings. We used the amplitude envelope of speech as the reference signal to investigate cortical alignment at the syllable rate. We segmented each participant’s MRI and then constructed the source space based on a warped Montreal Neurological Institute (MNI) anatomical grid template, which we used for group analysis. After preprocessing the individual MEG recordings, we computed the source modelling and cortico-acoustic coupling using the Dynamical Imaging of Coherent Sources (DICS) beamformer (41). Lastly, for statistical analysis we applied non-parametric randomization, cluster-based permutation statistical tests across participants for each frequency of interest and condition in five bilateral regions-of-interest (ROIs). We achieved the statistical assessment of cortico-acoustic coherence by comparison to two control conditions: coherence obtained either using surrogate data (trial shuffling) or using pre-stimulus data.

**Fig 2.**
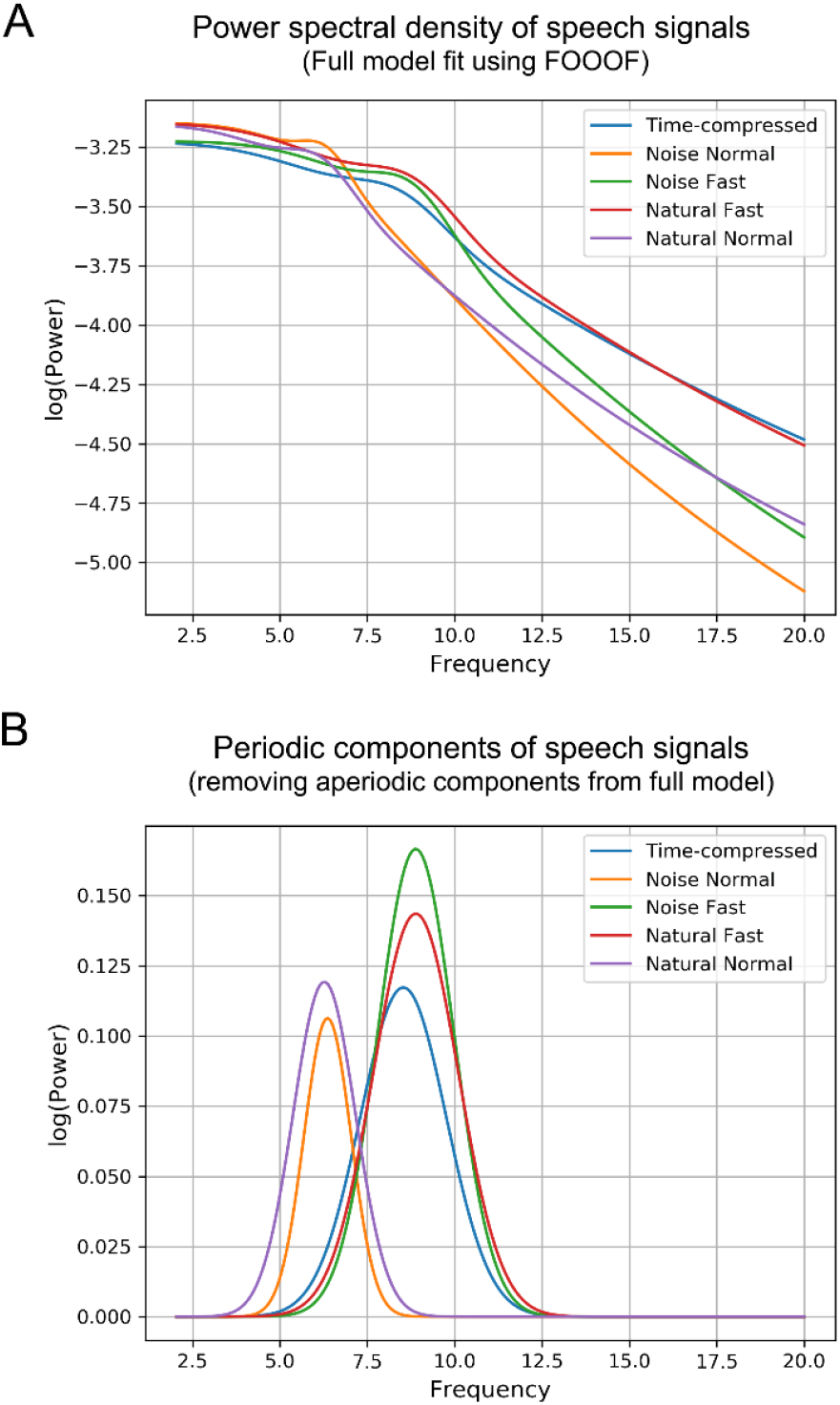
Power spectra of the acoustic signal for the five conditions. (A) Power spectra of the acoustic signals’ envelopes across the five experimental conditions: Natural Normal and Natural Fast correspond to naturally-produced speech at a normal (mean syllable rate 6.76 syllables/s) and fast rate (mean rate 9.15 syll/s), respectively. Time-compressed speech was compressed at the same syllable rate as in the natural fast rate condition (mean syllable rate 9.15 syll/s). Noise Normal and Noise Fast correspond to noise stimuli modulated with the amplitude envelopes of normal rate and fast rate sentences, respectively. The depicted spectra represent parametric model fits of the PSD (Power Spectral Density), that consist of aperiodic and periodic components computed using the FOOOF algorithm (42) (B) Periodic components of the full model shown in (A), i.e. after removal of the aperiodic (the so-called 1/f) component of the spectra. The spectral power peaks in the speech stimuli for each condition occur at frequencies that match the corresponding mean syllable rates calculated using the Praat software (see supplementary Table S1 and supplementary Fig S1 for details).

The ROI-based analysis using shuffled data as control revealed a shift in the frequency domain of cortical oscillations to align to the syllable rate of naturally-produced heard sentences. Significant increase of cortico-acoustic coherence for the normal rate condition was found at the frequency matching the mean normal syllable rate (∼6.25 Hz) (Fig. 3A). Crucially, for natural fast speech, coherence significantly increased at the frequency matching the mean fast syllable rate (∼8.75 Hz; Fig. 3B). The time-compressed speech condition did not show any significant brain coupling at the corresponding frequency (∼8.75 Hz). Comparable patterns of cortico-acoustic coupling were obtained when coherence during speech encoding was compared to coherence computed for baseline data (Fig. S2). As Fig. 3 shows, in both natural speech rate conditions, cortical tracking of speech envelope at the corresponding frequencies was seen in the primary auditory cortex (Brodmann Area BA 41), middle and superior temporal gyri (BAs 21/22), primary sensory (BA 1) and primary motor and premotor cortices (BAs 4/6) of the right hemisphere (see also Fig. S2). Note that we also found significant brain coupling in the right dorsal precentral gyrus (BA 4) for naturally and artificially accelerated speech at ∼6.25 Hz (normal rate frequency). By contrast, we did not find any significant increase in coherence for normal rate speech at the higher frequency (∼8.75 Hz). Neither of the two amplitude-modulated noise conditions presented statistically significant cortico-acoustic coupling at any of the two frequencies of interest (Figs. 3A and 2B).

**Fig. 3.**
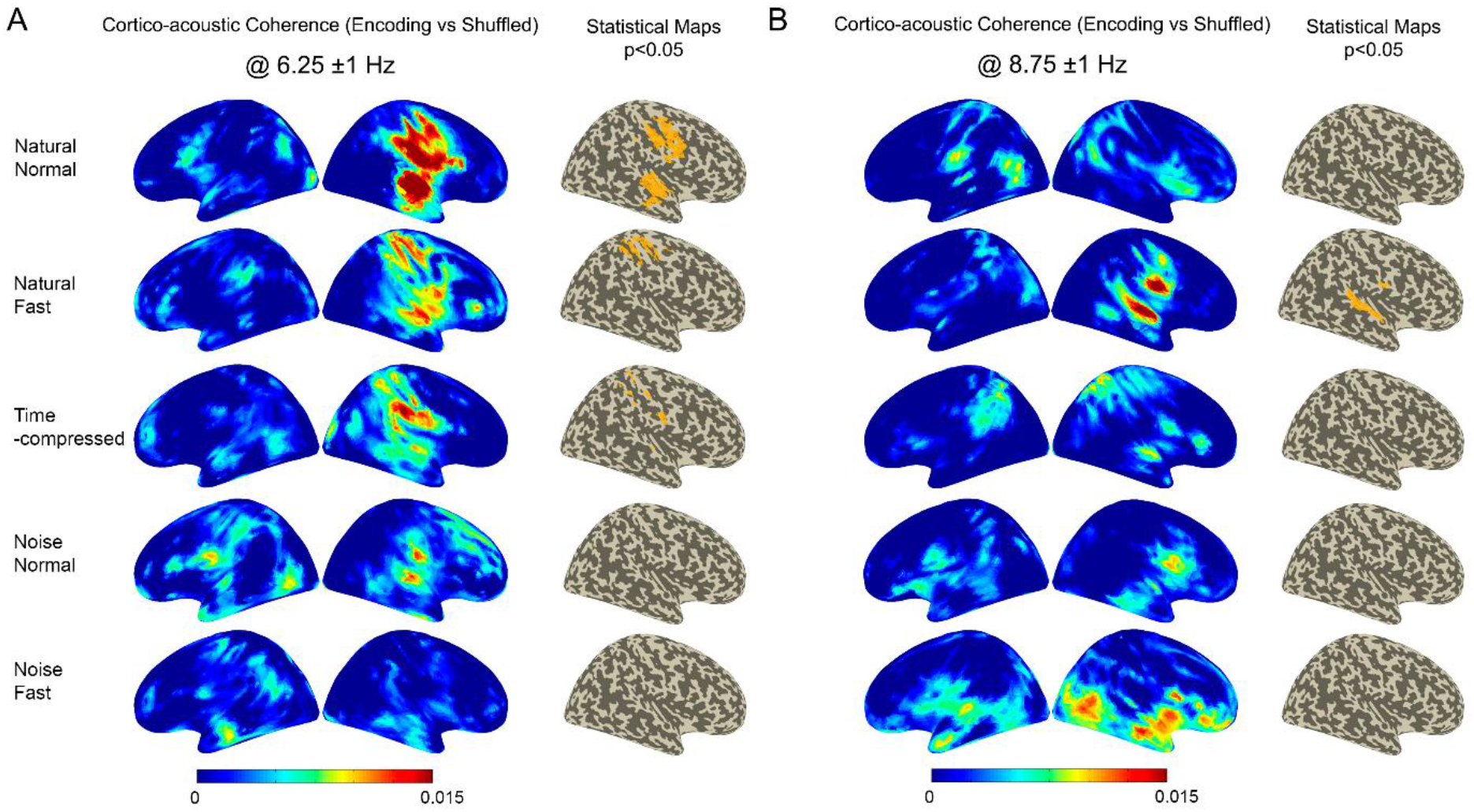
Cortical tracking of speech at (A) 6.25 (±1 Hz) and (B) 8.75 (±1 Hz) matching the normal and fast speech rates respectively. Coherence maps between signal’s amplitude envelope and neural oscillations in the active period (i.e. during stimulus presentation), as compared to surrogate data, together with statistical maps in the right hemisphere (corrected, α = .05; results were not significant in the left hemisphere) presented for the five conditions. Natural Normal = naturally-produced normal rate speech; Natural Fast = naturally-produced fast rate speech; Time-compressed = artificially accelerated speech from normal rate sentences at the same rate as natural fast speech; Noise Normal = amplitude-modulated noise with the envelope of normal rate sentences; Noise Fast = amplitude-modulated noise with the envelope of natural fast rate sentences.

To specifically test our hypothesis of a stronger involvement of the motor cortex in the tracking of natural fast as compared to time-compressed speech, we computed direct contrasts between speech conditions at the frequency matching the mean fast syllable rate (∼8.75 Hz). More specifically, we focused on a putative right articulatory area which we defined as the region of the right motor cortex (BA 4) that showed a cortico-acoustic coherence peak at ∼8.75 Hz (Fig. 3B) and based on previous neuroimaging reports on the involvement of the articulatory cortex in speech perception and production (43,44). Fig. 4A shows the maps of cortico-acoustic coherence for the three computed contrasts. Remarkably, the right articulatory motor cortex showed stronger entrainment to natural fast rate speech, compared to both time-compressed and natural normal speech (Fig. 4B, articulatory ROI). Note that contrasts between speech conditions using the whole set of predefined ROIs (as in Fig. 1) showed an increase of coherence for natural fast as compared to time-compressed speech in the same region as well as in the right auditory and temporal cortex (uncorrected results; see suppl Fig. S3). For completeness, we also computed direct pairwise contrasts between speech conditions at ∼6.25 Hz (normal syllable rate range). Results showed significantly stronger coupling to natural speech, both at normal and fast rates, compared to time-compressed speech in the right precentral ROI (see Fig. S4). The two naturally-produced conditions did not significantly differ from each other at the frequency of normal speech (∼6.25 Hz).

**Fig. 4.**
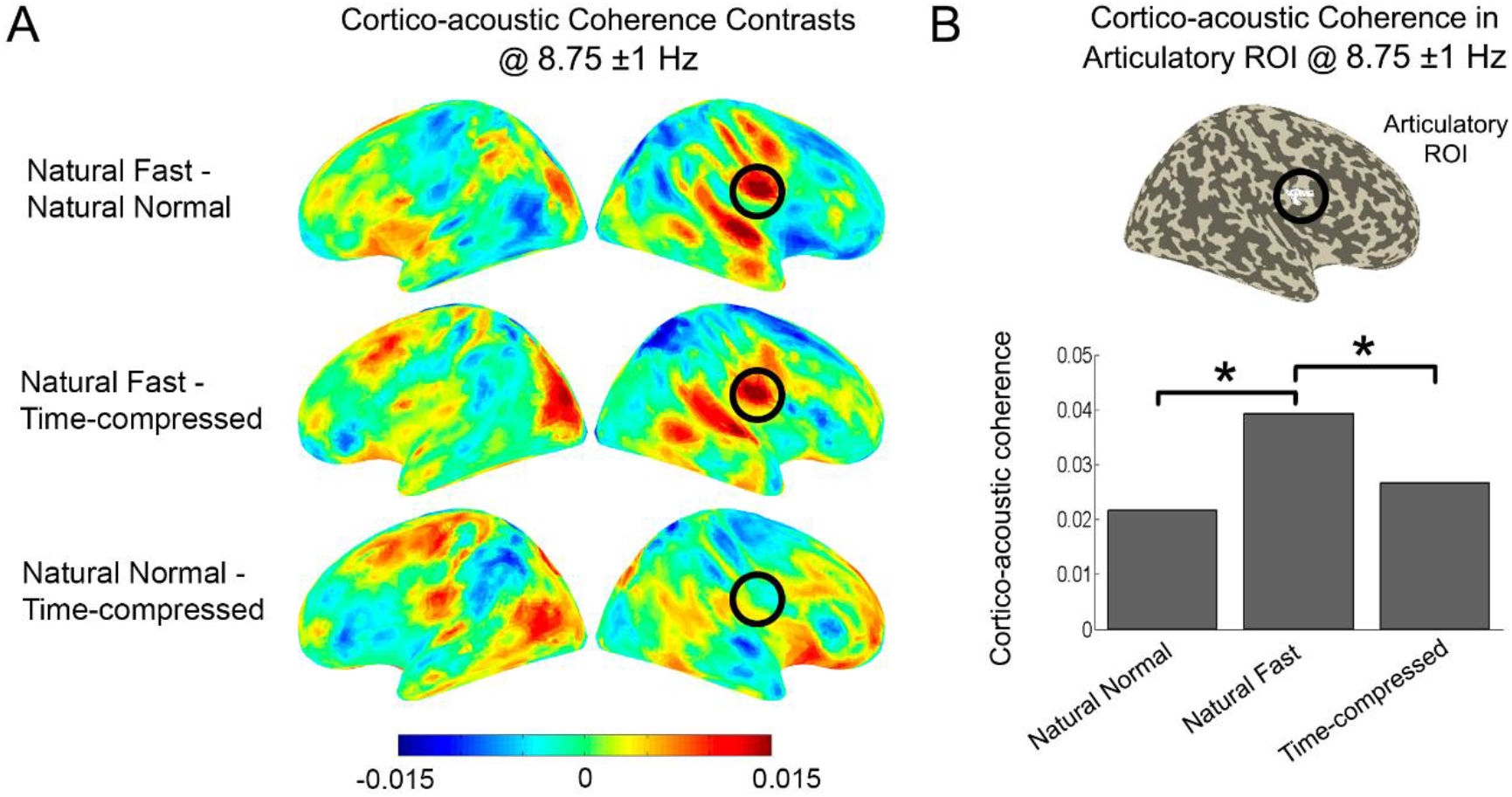
Direct contrasts between speech conditions at 8.75 (±1 Hz) reveal increased coupling to natural fast speech in articulatory cortex. (A) Contrast maps show the difference in cortico-acoustic coherence in the three pairs of speech conditions. The values represent the difference in Fisher z-transformed coherence between each two conditions. The black rings highlight the right articulatory ROI. (B) Mean cortico-acoustic coherence in the right articulatory ROI for the natural normal, natural fast and time-compressed conditions. Stars (*) indicate significant differences between conditions (t-test, α = .05, corrected).

Next, in a supplementary follow-up analysis, we sought to assess differences in cortico-cortical network dynamics associated with processing naturally and artificially accelerated speech. To this end, we measured seed-based phase interaction patterns at ∼8.75 Hz using weighted Phase Lag Index (wPLI). In light of the cortico-acoustic coherence results, we chose to focus the inter-areal coupling analyses on two key ROIs: the right articulatory motor and right auditory areas, as both showed stronger entrainment to natural fast than to time-compressed speech (Figs. 3B, 4 and S3). As illustrated in Fig. S6B, the right articulatory motor cortex showed higher inter-areal phase coupling for natural fast than for time-compressed sentences, mainly with the right auditory cortex (BA 41/42) and a left-lateralized network encompassing the inferior parietal cortex (supramarginal and angular gyri BA 39/40), Broca’s area (BA 44/45), the primary motor and premotor cortices (BAs 4/6) extending to the supplementary motor area (SMA), the primary sensory cortex (BA 1) and the dorsolateral prefrontal cortex (BA 9/10/46). Direct comparison between natural fast and normal rate speech at the same frequency (∼8.75 Hz) showed enhanced coupling between the right motor cortex and the left lateral premotor cortex (BA 6 including SMA) and left dorsolateral prefrontal cortex (BA 46).

When the right auditory cortex was used as the seed (Fig. S6A), results showed stronger coupling for the natural fast than for the time-compressed condition mainly with the left inferior parietal cortex (BA 39/40), Broca’s area, left inferior part of the primary motor and premotor cortices (BA 4/6), left primary sensory cortex (BA 1) and left primary auditory cortex (BA 41). For the contrast between natural fast and normal rate sentences, the right auditory cortex was more strongly coupled mainly to the bilateral primary motor and premotor cortices (BA 4/6, more ventrally in the right hemisphere) and right inferior frontal gyrus (BAs 44/45).

Finally, we sought to rule out that the observed increases in coupling between cortical oscillations and the speech envelope can be linked to increases in cortical power at the coupling frequency. To this end, we analyzed source power modulations, contrasting power for the speech encoding period with power measured during a pre-stimulus baseline. The analyses showed significant increases of spectral power at ∼6.25 Hz in the left prefrontal and inferior frontal cortex for all sentence conditions (Fig. S6). These areas did not overlap with the regions that exhibited significant cortico-acoustic coherence at this frequency. This was accompanied by significant desynchronization at ∼8.75 Hz in the right temporal cortex in all conditions (except for noise modulated with the envelope of fast rate sentences) and in the left inferior frontal cortex for the two naturally-produced speech conditions (Fig. S7). These power reductions in the same areas and frequencies in which we report entrainment to the natural fast stimuli suggest that the observed coupling cannot be attributed to local increases in power.

## Discussion

To date, most of the evidence for brain alignment to speech rate variations has come from research using artificially accelerated speech and has thereby left the issue of natural speech rate changes largely unaddressed. The present MEG study is the first to directly compare brain-to-speech coupling between naturally-produced fast speech and artificially compressed speech. We first showed that neural oscillations in auditory and (pre)motor cortex track natural variations of syllable rate at frequencies that specifically reflect the temporal structure of the speech material. Cortical rhythms indeed shift up their coupling frequency to match the faster modulations in natural fast speech. Surprisingly, we did not observe any significant cortical coupling at the same frequency (∼8.75 Hz) when the speech was generated through time-compression. Crucially, direct contrasts between conditions at this higher frequency revealed stronger tracking of speech envelope in the right motor cortex for natural fast than for artificially accelerated speech, possibly reflecting specific mapping to articulatory features of naturally-produced material. Furthermore, although the results of seed-based source-space wPLI analyses did not survive correction for multiple comparison across 8693 sources, the uncorrected statistics suggest that the right articulatory/motor ROI showed enhanced phase synchronization with the left temporo-parietal and motor cortices in the natural fast compared to the time-compressed condition. Finally, we found that the reported cortico-acoustic coupling is sensitive to the linguistic content of the stimuli as no significant increase of coherence is seen for amplitude-modulated noise, despite being generated using the speech signal envelopes.

### Auditory and motor cortices track syllabic rate changes in natural speech

Brain coupling to the amplitude envelope of naturally-produced speech was found in auditory and precentral regions of the right hemisphere, in agreement with oscillatory-based models of speech perception (4,6) and previous work showing right-lateralized brain responses to speech envelope (13,19,45–47). Specific MEG responses in the right pre/postcentral gyri were also recently reported for sequences of random syllables at 4 Hz (48). Most importantly, our coherence measures revealed that neural oscillatory activity is tuned to speech rate variations: cortical tracking of naturally-produced sentences was observed at frequencies coinciding with the syllable rate of the stimuli. For normal rate speech, cortico-acoustic coherence in auditory and (pre)motor regions increased at ∼6.25 Hz, whereas when participants listened to natural fast speech, the peak of coupling shifted up to ∼8.75 Hz (Fig. 3). Note that a significant increase of coherence in the right precentral cortex was also observed for naturally accelerated speech at the lower frequency. Syllable frequencies in natural speech tend to overlap between speech rates (49). Along this line, the envelope of our natural fast sentences also contains slower frequency components, which may account for the observed pattern of results. In fact, this explanation is consistent with the spectral power density plots (see supplementary Fig. S1). The lack of coupling for normal rate speech at the higher frequency (∼8.75 Hz), which was expected, however underlines the specificity of the reported effects. Similar patterns of brain-to-speech coupling were observed when we contrasted coherence between speech and MEG brain activity with coherence computed for baseline trials (supplementary Fig. S2). This replication using two distinct methods supports the reliability of our observations.

Importantly, the significant increases of cortico-acoustic coherence for natural normal and fast rate speech were not accompanied by power increases (but rather power suppression at ∼8.75 Hz) in the same cortical regions and frequencies (Figs S5 and S6), suggesting a genuine synchronization phenomenon that cannot be attributed to increases in signal amplitude. The power increase at ∼6.25 Hz (∼theta) to sentences in left prefrontal and inferior frontal cortex may be related to increased working memory load and lexico-semantic retrieval during sentence processing (50,51). Besides, the ∼8.75 Hz desynchronization in right anterior temporal and left inferior frontal regions agrees with studies showing alpha band (8-13 Hz) desynchronization during auditory stimulus processing. This is classically thought to reflect enhanced mental operations and thus more active cognitive processing of the signal (52,53). Alpha suppression associated with theta power enhancement in frontal regions have also been reported in response to speech for lexico-semantic processing (52,54,55).

Our findings first add new evidence to previous work on artificially accelerated speech (28,27) by revealing that cortical oscillations align to envelope modulations at a higher frequency to match the faster syllable rate of *naturally-produced* speech, despite increased articulatory variation (as compared to time-compressed speech). Such auditory and motor coupling at ∼8.75 Hz fits with recent results showing three peaks of resting-state theta-band activity in auditory cortex (4.5, 6.5 and 8.5 Hz) as well as intrinsic alpha-band activity (7-13 Hz) in the right precentral gyrus (56). It is also of note that the shift in coupling frequency was found for single, relatively short sentences with a syllable rate up to 10 syllables/s (mean = 9.15 syllables/s). Ahissar et al. (23) suggested that coupling to short sentences compressed to ratios of 0.35 (∼9 Hz) and 0.2 (∼14 Hz) failed in their experiment because neural oscillations may not have had enough time to change their coupling frequency to match that of the stimuli. Although this is a plausible explanation, the present data using naturally-produced material show that even with single and relatively short sentences, neural oscillations are able to adjust their coupling frequency to higher syllable rates.

In line with the study by Keitel and collaborators (39), our results emphasize the relevance of assessing neural tracking of speech at frequencies given by stimulus properties rather than in generic frequency bands which may not capture the specific underlying processes at stake. The MEG study by Alexandrou et al. (35) showed cortical tracking of spontaneously-produced connected speech at slow (∼2.6 syllables/s), normal (∼4.7 syllables/s) and fast (∼6.8 syllables/s) syllable production frequencies (yet slower than our fast rate condition). Despite providing valuable evidence regarding natural speech perception, the authors however examined coupling in the canonical delta (2-4 Hz) and theta (4-7 Hz) bands and did not look at potential variations in brain coupling frequency according to speech rates. Although our study used single sentences, we bring novel evidence for cortical alignment to natural syllable rates up to an average of 9 Hz and at frequencies that are specific to the speech material. Future work should certainly investigate brain coupling to longer extracts of naturally-produced speech at such normal and fast rates and in stimulus-based frequency bins as we did in our study.

### Role of motor cortex in tracking naturally produced fast speech

Assaneo and colleagues reported cortico-acoustic coupling in inferior and middle frontal gyri at 4.5 Hz (57), as well as enhanced phase coupling between both left and right auditory and motor regions (58), when participants listened to synthesized syllables at the same rate (see also 41 for delta motor coupling to normal rate sentences embedded in noise). Sheng and colleagues (48) also found specific tracking of syllables at a rate of 4 Hz in the right pre/postcentral cortex. Here, we show that right (pre)motor regions synchronize their oscillatory activity to more complex speech stimuli (i.e. meaningful sentences) that are naturally produced at faster rates (up to 9.15 syllables/s on average). Similar to auditory cortex, oscillations in the motor cortex are therefore able to shift up their coupling frequency to follow the natural increase in syllable rate. Remarkably, analyses at ∼8.75 Hz revealed tracking of natural fast speech in a region of the ventral motor cortex (BA 4) which coordinates are very close to those of the articulatory cortex (mouth motor region) identified in neuroimaging studies on speech production and/or perception (43,44,59). Coupling to naturally-produced fast speech in this ventral motor region may therefore reflect articulatory mechanisms and more particularly simulation of the syllable production rhythm of heard sentences.

Motor regions have also been suggested to contribute to top-down auditory processing and to the establishment of auditory temporal predictions (9,60–63). Our findings, along with those of few other studies (35,48,57), underline that motor regions do not only exert a modulatory control but directly track the speech signal. Neural oscillations in motor regions align to low-frequency modulations in natural speech, both at normal and fast rates, possibly reflecting sensorimotor integration processes. In the present study, participants listened to sentences with the same syntactic structure, it is therefore possible that motor regions tracked and synchronized to syllable rate regularities so as to predict the occurrence of the next syllables. Such a predictive timing mechanism may facilitate the syllabic parsing of the unfolding speech stream by the auditory cortex (61).

### Stronger motor coupling to naturally-produced than to artificially accelerated speech

No significant cortical coupling to time-compressed sentences was observed at the corresponding frequency (∼8.75 Hz), which is at odds with previous work, at least regarding auditory cortex oscillations (20,23,27). To the best of our knowledge, these studies have mostly focused on neural coupling in the auditory cortex and we are only aware of a few studies that documented theta (4-7 Hz) synchronization to degraded (though noise-vocoded, not time-compressed) speech in distributed cortical networks including the motor cortex (10,64). Note from Fig. 3B that coherence maps tend to show increase of coherence, although weaker, for time-compressed speech at ∼8.75 Hz in a similar fronto-temporal network as for natural fast rate speech, however this did not survive statistical correction despite a sample size of 23 analyzed participants. Some methodological considerations may account, at least partly, for this apparent discrepancy between studies. Unlike previous work, we mixed two types of accelerated speech, with sentences pseudo-randomly presented to participants. This may have elicited different coupling effects because of the attention-grasping nature of natural fast speech, which may be more difficult to process than time-compressed speech (30,32). Along this line, the increase of coherence at ∼8.75 Hz in the right temporal cortex for natural fast speech only (see Fig. 3B) may reflect encoding of greater spectro-temporal variations in this natural condition than in time-compressed speech, in line with the role of the right superior temporal gyrus in spectral processing (65–67). In fact, this stronger neural coupling to sentences in the natural fast compared to the time-compressed conditions in the right temporal cortex was also visible on the direct contrast map at the same frequency (∼8.75 Hz; see supplementary Fig. S3).

Most importantly, the way speech rate was sped up in our study may have affected brain oscillatory responses to sentences. Uttering speech at a fast rate is a nonlinear phenomenon whereby segments are not reduced similarly, partly because of articulatory constraints, thus enhancing the prosodic pattern (30,31). By contrast, artificially accelerated speech was obtained by linear compression, meaning that all segments were shortened in the same way. This leads to unnatural patterns that are not biologically (articulatory-speaking) plausible and may thus not resonate in brain motor regions (or less so) as naturally-accelerated speech does. Whereas we are indeed rather accustomed to and can reproduce natural fast sentences relatively easily, this is not the case for linearly time-compressed speech. Crucially, our results revealed significant coupling of the right motor cortex at ∼8.75 Hz to naturally accelerated but not to time-compressed speech, despite having the same syllable rate. Direct contrasts between the two conditions corroborated this result (Fig. 4). This major finding may reflect differences in the rhythmic structure of the two types of signals and demonstrate specific mapping, in the motor cortex, to articulatory features that characterize naturally-produced fast speech as compared to synthesized fast speech.

Remarkably, our contrasts analyses at the lower frequency (∼6.25 Hz) highlighted significantly stronger tuning in the precentral cortex for naturally-produced speech, either at a normal or fast rate, than for time-compressed speech (supplementary Fig. S4). No difference was observed when we contrasted the natural normal and fast rate conditions. This finding is particularly interesting and can be interpreted in the framework of studies showing that the motor cortex intrinsically oscillates in the theta band (56,68–70). Given that our ∼6.25 Hz frequency of interest fell within this range, our results may emphasize the tendency of the motor cortex to preferentially align to natural rather than to artificially speech. Hence, the motor cortex more strongly resonates, at its own preferred rhythm, with speech perception when the signal is naturally produced, irrespective of the syllable rate, than when it has been artificially manipulated. This may indicate enhanced synchronization to articulatory patterns that are most prominent in natural vs compressed speech as well as increased sensorimotor integration required to match phonological and articulatory templates of natural speech.

### Enhanced phase synchronization between right motor cortex and left dorsal stream for naturally-produced speech

Supplementary cortico-cortical coupling analyses at ∼8.75 Hz revealed enhanced connectivity between the right motor/articulatory cortex and the left inferior parietal and (pre)motor cortices and Broca’s area during perception of natural fast with respect to time-compressed sentences (supplementary Fig. S5). We also found the right auditory cortex to be more strongly coupled to these same regions in the naturally accelerated condition. This left-lateralized temporo-parieto-frontal network is part of the sensorimotor dorsal stream thought to instantiate forward and inverse articulatory-orosensory-auditory internal models to facilitate speech perception, especially under challenging conditions (36,37,71–77). In line with this, we show that listening to naturally-produced fast speech specifically increases the functional connectivity between the right motor cortex, which synchronizes to syllable rate, and regions of the left dorsal stream (extending to the SMA). Interestingly, such inter-hemispheric coupling was also described during the perception of distorted speech in a recent dual-coil TMS study (78). The results showed that disrupting the right ventral premotor cortex inhibited left motor cortex excitability, as reflected by decreased motor evoked potentials over lip muscles, when participants listened to imprecisely articulated syllables as compared to clear speech.

Consistent with the involvement of reverberant, dynamic bilateral speech motor networks in speech perception (79), our findings thus show that the right motor cortex specifically entrains to the syllable rate of naturally-accelerated speech, and is more strongly coupled to left parietal and (pre)motor regions when perceiving natural fast as compared to time-compressed speech. This may reflect enhanced resonance to the increased articulatory complexity of natural fast speech, which may more strongly rely on internal models than artificially accelerated speech to efficiently map distorted perceptually representations to stored orosensory and articulatory representations. Of course, one should keep in mind that the cortico-cortical phase synchronization patterns observed in the present study (Fig. S5) need to be interpreted with caution as the effects did not survive correction for multiple comparisons across the 8693 sources.

### Linguistic information influences cortico-acoustic coupling

Previous work documented auditory cortex alignment to both verbal and non-verbal stimuli (e.g., 44,79). Our data show that amplitude-modulated noise did not significantly “entrain” cortical oscillations at the two frequencies of interest, although these stimuli were generated using the temporal envelopes of normal rate and fast rate sentences. In other words, increased cortico-acoustic coherence was found for normal rate and fast rate sentences but not for stimuli with the same envelope characteristics but which lack linguistic information. This result suggests that brain coupling to natural syllable rate variations does not only reflect passive tracking of acoustic features present in the low modulations of the amplitude envelope, but that it is sensitive to the linguistic content of the heard items, consistent with previous studies (10,13,81–83). Alternatively, stronger coherence for sentences than for amplitude-modulated noise could result from the richer spectral structure of the former stimuli and not from the availability of linguistic information. Our results cannot currently disentangle these two interpretations and future work comparing brain oscillatory responses to unintelligible natural speech that has the same spectro-temporal complexity as intelligible natural speech are certainly needed (64).

## Conclusions

Our findings shed new light onto the brain oscillatory dynamics that mediate natural speech perception, by revealing that neural oscillations are tuned to natural speech rate variations at frequencies that match the syllabic structure of the acoustic input. We found such frequency-specific coupling not only in auditory but also in (pre)motor regions, emphasizing their role in speech sensorimotor integration. Our results also provide unprecedented evidence for a stronger oscillatory coupling in right motor cortex to naturally accelerated compared to artificially manipulated speech. In addition, our follow-up cortico-cortical connectivity analysis suggests enhanced coupling between right motor cortex and left parietal and motor regions for natural fast speech. These observations likely reflect enhanced distributed tracking and encoding of articulatory features of naturally accelerated speech. Our data thus highlight the relevance of using both natural speech material (despite being more methodologically constraining) and stimulus-specific (vs generic) frequencies to thoroughly assess brain-to-speech alignment in future studies. The prominent role of right auditory but also motor areas unveiled by our study might provide valuable insights for the advancement of oscillatory models of speech perception and production Finally, the proposed paradigm may also prove of high interest to investigate the developmental trajectory of neural tracking of speech, both in children with typical and atypical language development.

## Materials and Methods

### Participants

Twenty-four French native speakers participated in the study after providing informed consent (14 females, mean age 23 years, range 18-45 years). All participants were right-handed (mean score at the Edinburgh handedness inventory = 94) (84) and reported normal hearing together with no history of neurological or psychiatric disorder. The protocol conformed to the Declaration of Helsinki and was approved by the local ethical committee (Comité de Protection des Personnes Lyon Sud-Est II; ID RCB: 2012-A00857-36). Participants received monetary compensation for their participation.

### Stimuli

We created 288 meaningful sentences (7-9 words) following the same syntactic structure: Determiner – Noun 1 – Verb – Determiner – Noun 2 – Preposition – Determiner – Noun 3 (e.g., “Sa fille déteste la nourriture de la cantine” / *His daughter hates the food at the canteen*). Sentences were recorded in a sound-attenuated booth by a French native professional theatre actor who was able to produce sentences at the required fast rate while remaining intelligible. We recorded each sentence twice (44.1 kHz, mono, 16 bits) using ROC*me*! software (85), first at a normal and then at a fast rate. The procedure was the following: the sentence was first displayed on a computer screen in front of the speaker who was instructed to silently read it and to subsequently produce it aloud as a declarative statement at his normal speech rate. The speaker then produced the sentences at a fast rate (i.e. as fast as possible while remaining intelligible) using the same procedure (i.e. no external pacing was imposed).

We calculated the durations of the 2×288 sentences and the number of actually produced syllables for each sentence with Praat software (40). The mean syllable rate was 6.76 syllables/s (SD 0.57) for natural normal rate sentences and 9.15 syllables/s (SD 0.60) for natural fast rate sentences. This led to an overall fast-to-normal ratio of 0.74 (i.e. speed-up factor of 1.35). Subsequently, we computed time-compressed sentences by digitally shortening them with a PSOLA (Pitch Synchronous Overlap and Add) algorithm (86), as implemented in Praat. We obtained compression rates for each sentence: we matched each individual time-compressed sentence in terms of syllable rate to its equivalent natural fast item. This artificial compression corresponds to a re-synthesis of the natural normal rate stimulus, changing only its temporal structure. For the total of 864 sound files (288×3 speech rate variants), we applied an 80 Hz high-pass filter, and smoothed the amplitude envelope sentence-initially and finally. We then peak normalized the intensity of the sound files.

Finally, for each of the 288 normal rate and 288 fast rate sentences, we created amplitude-modulated noise stimuli (i.e., Gaussian white noise with no linguistic content) using the amplitude envelope of the sentence material. These stimuli served as control non-speech conditions. We also used 48 filler sentences (different from the experimental stimuli but with the same syntactic structure, either at a normal rate, natural fast rate, or time-compressed) in which we added beep-sounds at the end and to which the participants had to respond during the experiment (see MEG data acquisition and task design).

We divided the total number of stimuli into two experimental lists, each including 288 sentences (96 normal rate, 96 natural fast rate and 96 time-compressed), 192 amplitude-modulated noise stimuli (96 with normal rate envelope and 96 with fast rate envelope) and 48 filler sentences. Each stimulus appeared in each rate condition across all participants but only once per list (to avoid repetition effects).

### MEG data acquisition and task design

Participants were comfortably sitting in a sound-attenuated, magnetically-shielded recording room with a screen in front of them. We presented stimuli binaurally through air-conducting tubes with foam insert earphones (Etymotic ER2 and ER3). Prior to the MEG recording, we determined participants’ auditory detection thresholds for each ear with a 1-minute pure tone of 44 kHz; the level was then adjusted so that we presented the stimuli at 50 dB Sensation Level with a central position (stereo) with respect to the participant’s head.

During the experiment, participants attentively listened to all stimuli from one of the two experimental lists while looking at a fixation cross at the center of the screen. Their task was to detect beep-sounds embedded in filler sentences by pressing a button (response button Neuroscan, Pantev) with their left index finger. Participants detected 100% of these trials, which we excluded from subsequent analysis. We pseudo-randomly presented all stimuli in 8 blocks of 66 trials allowing for short breaks. A training phase with 5 sentences (different from experimental stimuli) preceded the actual experiment. Each trial started with the appearance of a fixation cross which remained on the screen throughout the duration of the trial. An auditory stimulus (sentence or amplitude-modulated noise) was delivered 1500 ms after trial onset. Each trial was followed by an inter-trial interval (grey screen) of 1250 ms. We instructed the participants to attentively listen to the presented stimuli. In the case of filler stimuli, they had to press the response button as quickly as possible. To maintain the participants’ attention throughout the experiment, we informed them there would be questions about the content of the sentences at the end of the experiment. We used Presentation software (Neurobehavioral Systems) to run the experiment.

We recorded brain activity of the 24 participants using a 275-channel whole-head MEG system (CTF OMEGA 275, Canada) at 1200 Hz sampling rate. We placed three fiducial coils (nasion, left and right pre-auricular points) on each participant to determine head position within the MEG helmet. We also placed four electrooculographic (EOG) electrodes to record horizontal and vertical eye movements. We monitored reference head position before each of the 8 experimental blocks and tracked head movements throughout the experiment using continuous head position identification (HPI).

### Data analysis

We performed all analyses using custom written Matlab scripts (Mathworks Inc., MA, USA) and the Fieldtrip toolbox (87). Fig. 1 describes the general methodology. The following section describes the processing of (1) speech recordings, (2) MEG data, (3) Magnetic Resonance Imaging (MRI) data, (4) source localization and coherence analyses, and (5) statistical analyses.

#### Speech recordings (reference signals)

We computed the amplitude envelope of the speech signal following the methodology of Peelle and colleagues (10). We first rectified the signal (full wave rectification) and then filtered it using a Butterworth low-pass filter (30 Hz, fifth order Butterworth filter, zero-phase digital filtering). For the cortico-acoustic coupling analysis, we selected speech envelope segments between 200 and 1000 ms post-stimulus onset (equal to the time of interest of the functional MEG data). We computed the spectral power for all sentence envelopes for each condition and identified the central frequencies of the prominent rhythmic components, using the FOOOF algorithm (fitting a parameteric model to the power spectral densities PSD) and subtracting the 1/f (i.e. aperiodic) component (42). The main peaks in the speech PSD were found at 6.35 Hz for normal rate, 8.91 Hz for natural fast rate and 8.61 Hz for time-compressed speech (Fig. 2, see supplementary Table S1 and Fig. S1 for details), which closely match the mean syllable rates calculated with Praat (6.76 syllables/s (SD 0.57) for natural normal rate and 9.15 syllables/s (SD 0.60) for natural fast and time-compressed speech). The frequency peaks identified in the spectra of the speech signals using the FOOOF method were used to define the frequencies of interest for the coherence analysis, namely 6.25 (±1 Hz) and 8.75 (±1 Hz) for the normal and fast speech rates respectively.

#### MEG data

We first segmented the MEG data into periods of 3 s (from 1 s before stimulus onset to 2 s after onset) for preprocessing. We rejected data segments contaminated by eye blinks, heartbeat and muscle artefacts using a standard semi-automatic procedure available in the Fieldtrip toolbox as follows. First, we filtered the signals at 50, 100 and 150 Hz, we then re-sampled the data to 300 Hz and rejected the deviant trials from visual inspection. Second, we detected and rejected EOG artifacts, jumps and muscle artifacts and visually double-checked the trials. Finally, we performed an Independent Component Analysis (ICA) to correct for electrocardiographic (ECG) artifacts as well as to check for residual EOG artifacts. For each trial, we defined the speech encoding period (i.e. active period) as the time from 200 ms to 1000 ms post-stimulus onset, and the baseline from 1000 ms to 200 ms before onset.

#### MRI data

We acquired the T1-weighted structural MRIs (MRI 1.5 T, Siemens AvantoFit) of 23 of the 24 participants after the MEG study. We aligned each MRI to the MEG coordinate system using the localization coils marked on the MRIs and the interactive alignment option in the Fieldtrip toolbox. We segmented each MRI and then computed each subject’s head model and source space using Fieldtrip. We used the single-shell as volume conductor model (88). To have a common space for group analysis, we constructed each subject’s source space based on a warped MNI template grid. This way, each location on the template grid corresponds to homologous grid points across subjects and therefore, we can average results across subjects at the source level. We used a template grid with 8693 vertices.

#### Source localization, power and cortico-acoustic coherence estimation

We estimated source power and cortico-acoustic coupling using Dynamical Imaging of Coherent Sources (DICS) beamformer (41). This method allowed us to appropriately assess cortico-acoustic coupling by computing coherence between speech envelope and cortical activity.

The magnitude squared coherence is defined as the linear correlation between two signals as a function of frequency. It is mathematically defined by:

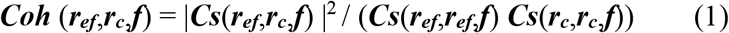

Where ***r***_***ef***_ is the reference signal, namely the amplitude envelope of the speech signal; ***r***_***c***_ is the signal at each vertex of the anatomical grid estimated with DICS; ***f*** is the frequency bin and ***Cs*** is the cross spectral density matrix.

We computed the cross spectral density (CSD) matrix using the multitaper FFT with Slepian tapers (i.e. Discrete Prolate Spheroidal Sequences, DPSS), at the frequency bins closest to the frequencies of interest, with ±1 Hz of spectral smoothing. The identification of the frequencies of interest for the normal and fast conditions was based on the FOOOF analysis. Given the length of the data windows (0.8s), the coherence frequency bins closest to the peaks identified from the spectra of the speech signals (using FOOOF) turned out to be 6.25 and 8.75 Hz for the normal and fast speech conditions respectively.

In addition to computing cortico-acoustic coherence during the speech encoding period (200 to 1000 ms), we also computed cortico-acoustic coherence (a) for the baseline window (−1000 to -200 ms) and (b) for shuffled data, as control conditions for statistical analysis. In the case of shuffled data, we randomly permuted the speech trial order (i.e. destroying the correspondence between the speech stimulus heard by the participant and the associated MEG data segment) before computing cortico-acoustic coherence. We repeated the shuffling procedure 100 times and averaged coherence across iterations for each participant, each condition, each node and frequency of interest. This led to 23 surrogate coherence values associated with 23 true coherence values, for each node, frequency and condition.

#### Cortico-cortical connectivity analyses

To quantify task-based modulations of cortico-cortical interaction patterns, we computed seed-based connectivity via weighted Phase Lag Index (wPLI). To this end, we used two key ROI as seeds: the right auditory ROI and the right articulatory/motor ROI. This choice was based on the results of the cortico-acoustic coupling analyses, as it was intended as an additional follow-up analysis. For each participant, we computed the wPLI connectivity matrix in the frequency band ∼8.75 +/-1 Hz for three distinct conditions: time-compressed, natural fast rate and normal rate speech. We chose wPLI because of its relative insensitivity to linear mixing effects compared to other metrics such as coherence or phase-locking value (89,90).

### Statistical analyses

We conducted group statistical analysis for the 23 participants (out of 24) with individual MRI. As control conditions, we used shuffled data and the baseline period for cortico-acoustic coherence, and the baseline for source power analysis. We compared the speech encoding period to each control condition applying non-parametric Monte-Carlo randomization (1000 randomizations), dependent-samples T-test statistics using FieldTrip. We corrected for multiple comparisons using ‘maxsum’ cluster-based correction and used a statistical significance threshold of 0.05 (91). Because we hypothesized that brain-speech coherence increases compared to control conditions, we used one-sided tests for the coherence analyses. However, two-sided tests were used when we explored task-based differences in spectral power, since we tested for both increases and decreases of mean power. Furthermore, and in line with several previous studies on neural entrainment to speech (10,28,60), statistical analyses were performed on regions of interest (ROIs). These were determined using the Automated Anatomical Labeling (AAL) atlas (92). We chose ten ROIs (five in each hemisphere) based on neurocognitive models (5,36) and neuroimaging data on speech perception (74,93,94). The ROIs consisted of Heschl’s gyrus, superior temporal gyrus, middle temporal gyrus, precentral and postcentral gyri (bilaterally). For the analysis of direct contrasts between speech conditions, we additionally defined the motor/articulatory cortex ROI as all the voxels located within 8 mm from two locations in MNI space at (66, 3, 17) and (66, -4, 17). These were chosen based respectively on (a) previous reports identifying articulatory cortex (43,44) and (b) the location of the speech-brain coherence peak found at ∼8.75 Hz. Finally, we used the BioImage Suite software (www.bioimagesuite.org) (95,96) to identify the Brodmann areas detected as significant in the statistical analysis. As for the wPLI analyses, we contrasted the natural fast with (a) the time-compressed condition and (b) the normal rate condition using non-parametric Monte-Carlo randomization (10000 randomizations) and dependent-samples one-sided T-test statistics using FieldTrip.

## Acknowledgements

We would like to thank Damien Gouy for recording the sentences, Emmanuel Ferragne for his help with Praat software, and Mélanie Canault and François Pellegrino for discussions on articulatory phonetics and signal processing respectively. We also thank Sébastien Daligault and Claude Delpuech from the CERMEP of Lyon for assisting in MEG data acquisition, and Mainak Jas for his implementation of the algorithm for automatic parameterization of neural power spectral densities (PSDs). V.B. was supported by funding from the French National Research Agency (ANR) for the ODYSSEE project (ANR-11-JSH2 005 1) and by the LabEx ASLAN (ANR-10-LABX-0081) of Université de Lyon within the program “Investissements d’Avenir” (ANR-11-IDEX-0007) of the French government operated by the ANR. K.J. is supported by funding from the Canada Research Chairs program and a Discovery Grant (RGPIN-2015-04854) from the Natural Sciences and Engineering Research Council of Canada, a New Investigators Award from the Fonds de Recherche du Québec - Nature et Technologies (2018-NC-206005) and an IVADO-Apogée fundamental research project grant.

