## Supplementrary Material for "Neural oscillations track natural but not artificial fast speech: Novel insights from speech-brain coupling using MEG"

SUPPLEMENTARY TABLES

**Table S1.** Centre frequencies (mean of the Gaussian) at which power spectrum peaks (maximum distance between the background aperiodic fit and the peak of the Gaussian in the power spectra model) were found for each experimental condition using the FOOOF algorithm which provides automatic parameterization of neural power spectral densities (PSDs) (43). Amplitude of maximal peaks (in arbitrary units, au), bandwidth of the Gaussian (in Hz, 2 standard deviations), as well as goodness of fit metrics ($R^2$ and Root Mean Square Error, RMSE) are reported.

| | Centre frequency (Hz) | Amplitude of maximal peak (au) | Bandwidth (Hz) | Goodness of fit |
|---|---|---|---|---|
| Natural Normal | 6.35 | 0.102 | 1.88 | $R^2$ = 0.999, RMSE = 0.011 |
| Noise Normal | 6.45 | 0.082 | 1.46 | $R^2$ = 0.999, RMSE = 0.014 |
| Natural Fast | 8.91 | 0.134 | 2.44 | $R^2$ = 0.996, RMSE = 0.021 |
| Time-Compressed | 8.61 | 0.111 | 2.39 | $R^2$ = 0.998, RMSE = 0.014 |
| Noise Fast | 8.95 | 0.147 | 2.23 | $R^2$ = 0.997, RMSE = 0.020 |

### SUPPLEMENTARY FIGURES

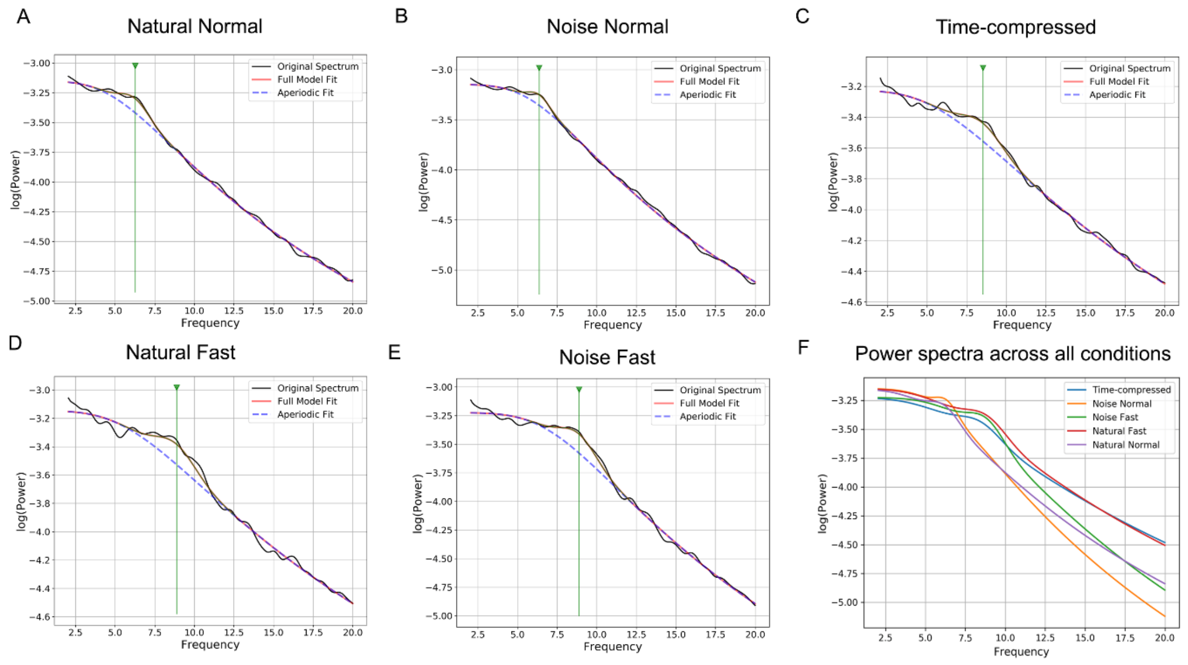


**Fig. S1.** Spectral power plots for the acoustic stimuli (speech and noise signals) for all five conditions (A-E), and an overview of the fitted model across all conditions (F). The model fitting (periodic + aperiodic components) was performed using the FOOOF algorithm (43), and all details are provided in Table S1.

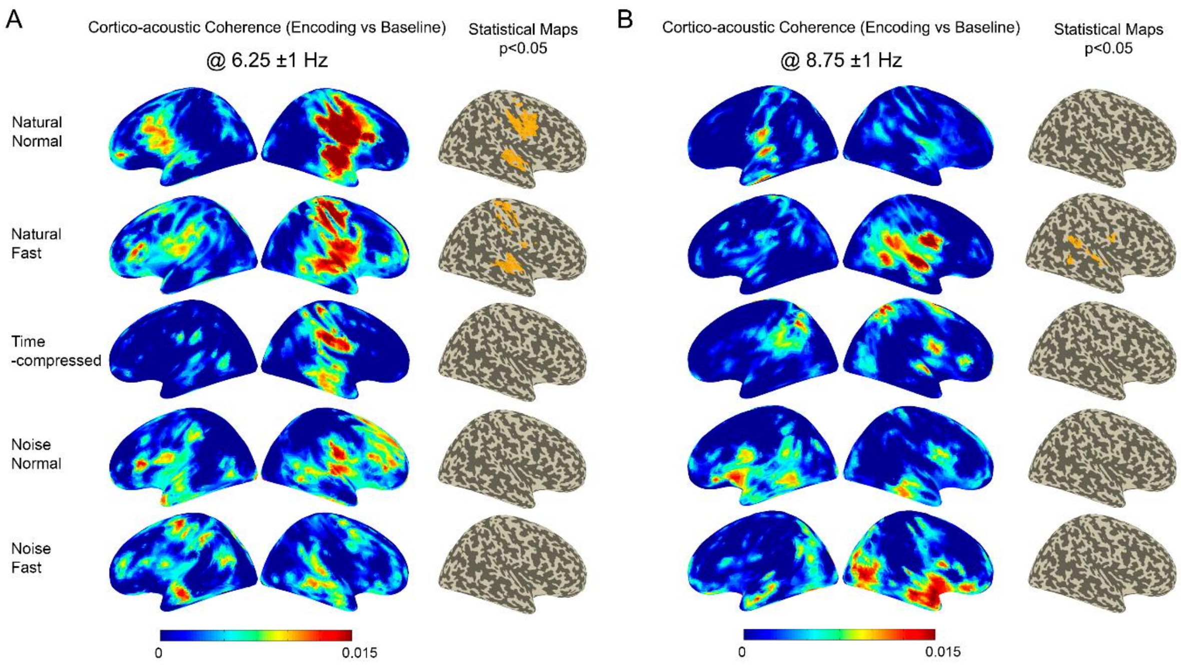


**Fig. S2. Cortical coupling to speech vs baseline data at (A) 6.25 (±1 Hz) and (B) 8.75 (±1 Hz) matching the normal and fast speech rates respectively.** Coherence maps between amplitude envelope of acoustic stimuli and MEG brain activity during stimulus presentation as compared with coherence computed for pre-stimulus baseline are presented for each condition. Statistical maps for the right hemisphere (corrected, $\alpha = .05$) are also shown (no significant increase of coherence was found in the left hemisphere). These results confirm the findings reported in Fig. 3, using a different control condition.

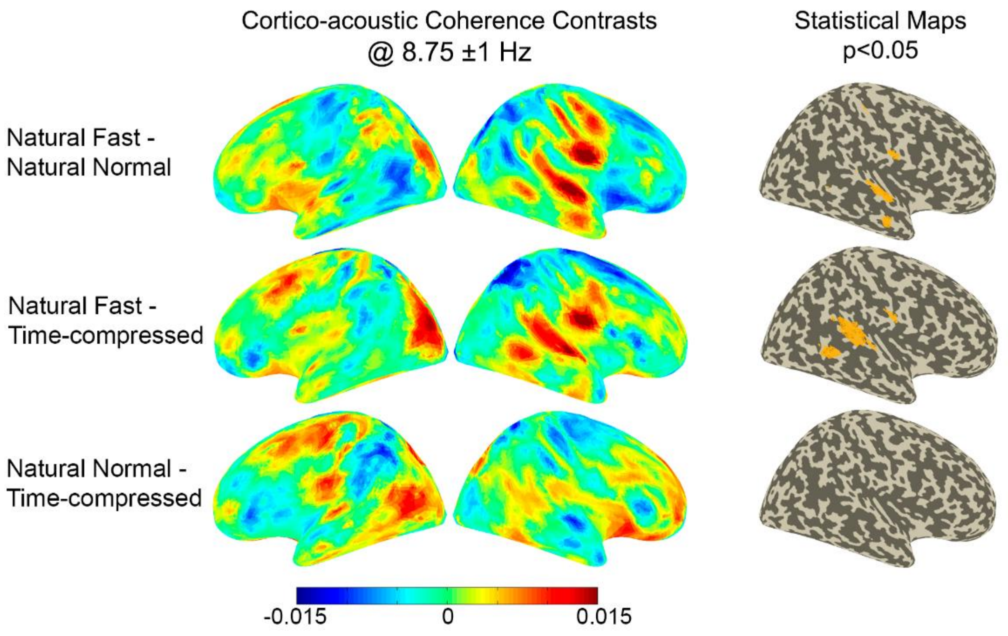


**Fig. S3. Direct contrasts between speech conditions at 8.75 (±1 Hz) in the set of predefined ROIs (cf. Fig. 1).** Maps of coherence computed between speech amplitude envelope and active period contrasted between the three speech conditions together with statistical maps for the right hemisphere (note that these are uncorrected maps, α = .05) are reported.

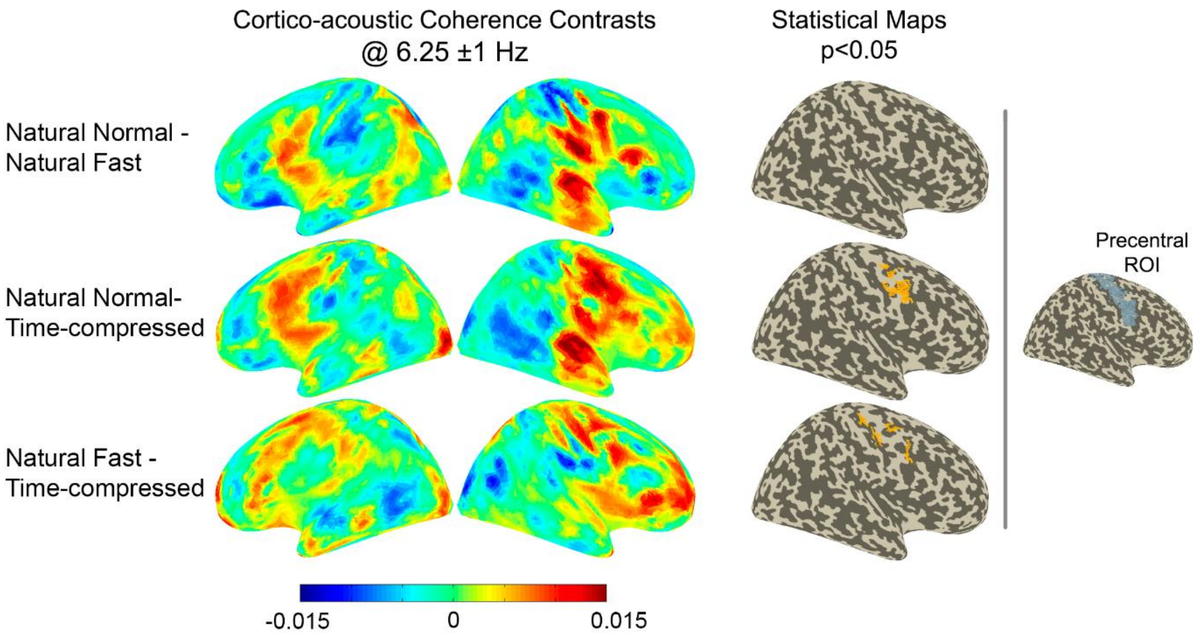


**Fig. S4. Direct contrasts between the three speech conditions at 6.25 (±1 Hz) in the right precentral ROI.** Maps of cortico-acoustic coherence contrasts as well as statistical maps for the right hemisphere (corrected, α = .05) are shown.

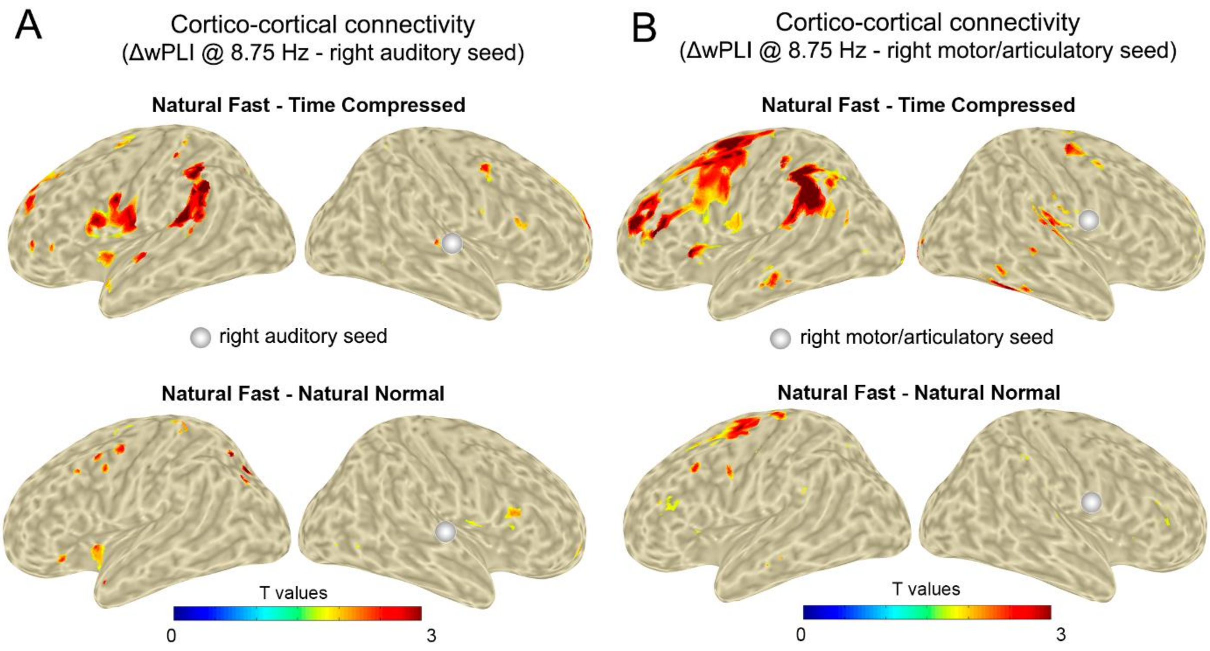


**Fig. S5. Task-based modulations of cortico-cortical phase interactions at 8.75 (±1 Hz) using seed-based wPLI coupling analyses. (A)** Contrasts between Natural Fast and Time-compressed conditions reveal areas of enhanced wPLI coupling with activity in a right auditory cortex seed (upper panel). By contrast, comparing the Natural Fast condition with the Natural Normal condition revealed less differences (lower panel). **(B)** Same as in panel (A) but for a right motor/articulatory seed. Note that the two reference seeds were chosen based on the previous cortico-acoustic analyses. The color bars indicate t-values based on Monte-Carlo randomizations (1000 permutations, one-tailed, uncorrected).

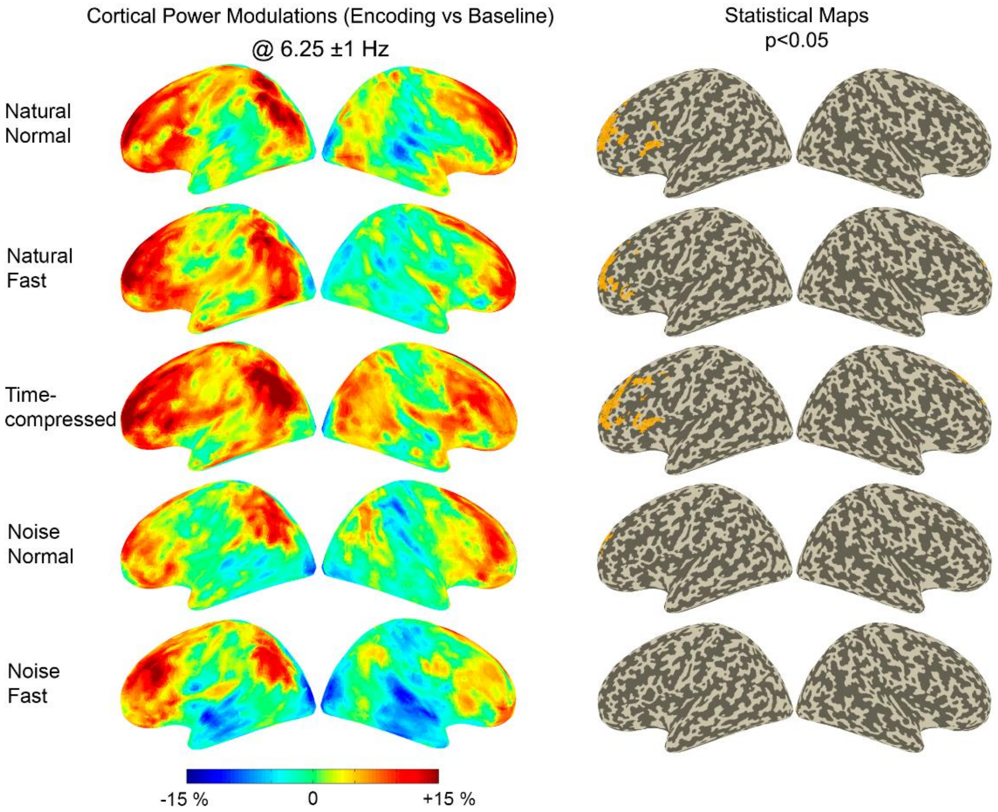


**Fig. S6. Power modulations at 6.25 (±1 Hz) (i.e. normal syllable rate).** Power maps and statistical maps (corrected, $\alpha = .05$) are reported for the speech and noise conditions as compared to pre-stimulus baseline.

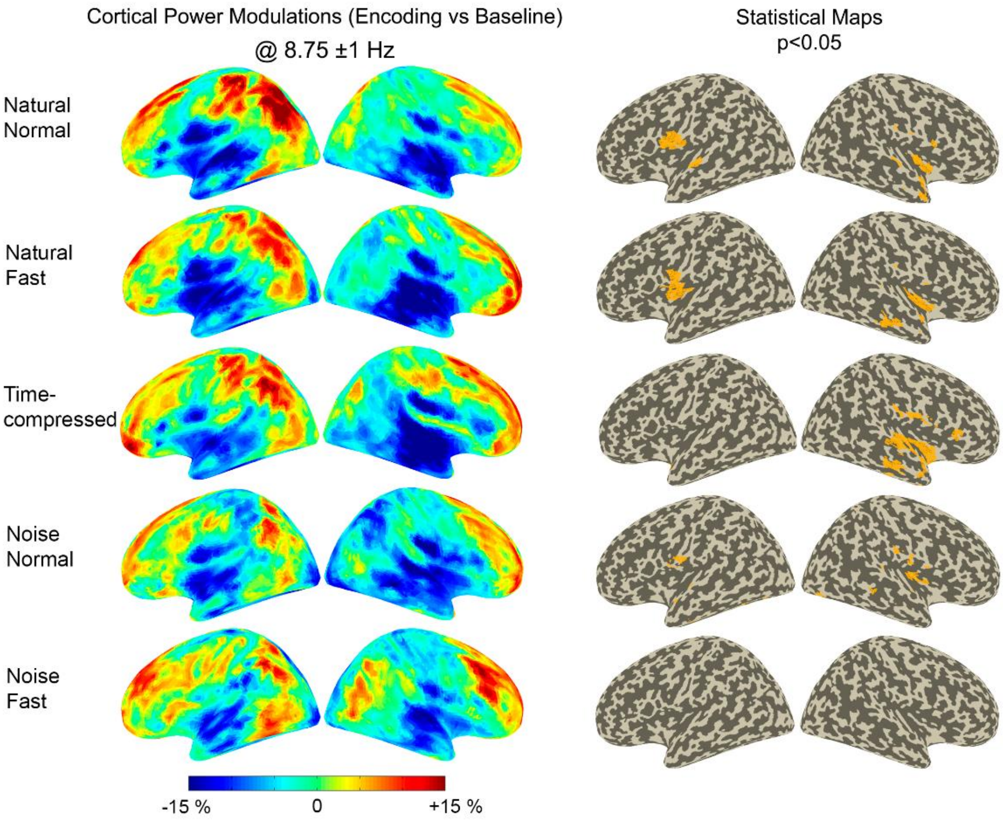


**Fig. S7. Power modulations at 8.75 (±1 Hz) (i.e. natural fast and time-compressed syllable rates).** Power maps and statistical maps (corrected, α = .05) for the different conditions as compared to baseline are shown.
